# The diverse viromes of Australian lizards are shaped by host taxonomy and habitat

**DOI:** 10.1101/2024.01.24.577151

**Authors:** Jackie E. Mahar, Michelle Wille, Erin Harvey, Craig C. Moritz, Edward C. Holmes

## Abstract

Lizards inhabit diverse ecologies and evolutionary histories and hence represent a promising group to explore how hosts shape virome structure and virus evolution. Yet little is known about the viromes of these animals. In Australia, squamates (lizards and snakes) comprise the most diverse order of vertebrates, and Australia hosts the highest diversity of lizards globally, with the greatest breadth of habitat use. We used meta-transcriptomic sequencing to determine the virome of nine co-distributed, tropical lizard species from three taxonomic families in Australia and analyzed these data to identify host traits associated with viral abundance and diversity. We show that lizards carry a large diversity of viruses, identifying more than 30 novel, highly divergent vertebrate-associated viruses. These viruses were from nine viral families, including several that contain well known pathogens, such as the *Flaviviridae*, *Picornaviridae*, *Bornaviridae, Iridoviridae* and *Rhabdoviridae*. Members of the *Flaviviridae* were particularly abundant across species sampled here, largely belonging to the genus *Hepacivirus*: 14 novel *Hepaciviruses* were identified, broadening the known diversity of this group and better defining its evolution by uncovering new reptilian clades. The evolutionary histories of the viruses studied here frequently aligned with the biogeographic and phylogenetic histories of the hosts, indicating that exogenous viruses may help infer host evolutionary history if sampling is strategic and sampling density high enough. Notably, analysis of alpha and beta diversity revealed that virome composition and richness was shaped by host taxonomy, habitat and range size. In sum, we identified a diverse range of reptile viruses that broadly contributes to our understanding of virus-host ecology and evolution.

## 1. Introduction

Animal virology has traditionally focused on viruses of mammals and birds as they are the most likely natural reservoir hosts for viruses that may emerge in humans and animals of economic importance (Mollentze and Streicker 2020; Zhang et al. 2018). While this has provided major insights, it has necessarily resulted in a skewed view of viral diversity that limits our understanding of virus evolution and ecology, including the frequency and determinants of cross-species transmission and host range. Indeed, many viral families traditionally associated with mammals and birds have now been found in other vertebrate classes such as amphibians, reptiles and fish (Harding et al. 2022; Shi et al. 2018). Since far greater biological diversity exists within these other vertebrates – that comprise ∼33,000 documented species – it is reasonable to assume that they also harbour a substantial diversity of viruses (Zhang et al. 2018).

A variety of factors make lizards particularly informative for the study of viral ecology and evolution. Squamates (lizards and snakes) are a highly diverse and successful group of vertebrates, comprising 96.3% of non-avian reptile diversity (Pincheira-Donoso et al. 2013), with over 10,000 extant species globally (Herrera-Flores et al. 2021). Squamates exhibit a range of morphologies, life history traits, and diverse ecologies, inhabiting every continent except Antarctica (Pincheira-Donoso et al. 2013; Pyron, Burbrink and Wiens 2013). Australia is home to the largest number of reptile species, comprising ∼10% of the world’s reptiles (Geyle et al. 2021), of which squamates are the most diverse vertebrate order (Wilson and Swan 2017). These animals are well adapted to the Australian landscape, colonising and radiating across a diverse range of environments and ecological niches (Pianka 1969; Pianka 1973; Morton and James 1988;). Australian lizards comprise old endemic Gondwanan lineages that pre-date the isolation of Australia and Antarctica, as well as more recent immigrant lineages from the North (Brennan and Oliver 2017). For example, the Gondwanan group of Pygopodoidea have a crown age of greater than 50 million years, while the genus *Gehyra* have a crown age in the mid-Miocene ∼20 million years ago, having colonized from Asia (Oliver and Hugall 2017). Several squamate species are at high risk of extinction within the next twenty years (Geyle et al. 2021) and infectious disease emergence can severely threaten reptile populations (Zhang et al. 2018).

A key, yet rarely addressed, aspect of virus evolution and emergence is understanding the traits that determine the diversity and abundance of viruses carried by a host (Wille et al. 2019). A number of studies have linked viral diversity and abundance to specific host traits, including phylogenetic history, habitat, body mass, geographic range, community diversity, biome, location, and infection status with particular pathogens (Olival et al. 2017; Wille et al. 2018; Geoghegan et al. 2021). However, the effects of host ecology on viral diversity have only been explored in a handful of systems with varying results. To better understand the diversity of viruses carried in lizards and how host traits affect viral abundance and diversity we used meta-transcriptomic sequencing to explore the virome of nine lizard species residing in various environments and habitats across the biologically diverse monsoonal tropics of northern Australia.

## 2. Methods

### 2.1 Ethics statement

All work was carried out according to the Australian Code for the Care and Use of Animals for Scientific Purposes with approval from the institutional animal ethics committee (Permit ANU animal ethics approval A2016-07) and State authorities (collecting permits NT 58454 and WA SF010911).

### 2.2 Animal Sampling

Liver samples were collected from apparently healthy *Carlia amax*, *Carlia munda*, *Carlia sexdentata*, *Cryptoblepharus metallicus*, *Heteronotia planiceps*, *Heteronotia binoei*, *Gehyra nana*, *Gehyra arnhemica*, and *Oedura marmorata*. Lizards were collected in autumn and winter months between April 2016 and June 2017, in arid regions of the eastern Kimberley and mesic regions of the “Top End”, Australia (Table 1, Table S1). Samples were collected in Australian bioregions Arnhem Coast (ARC), Victoria Bonaparte (VIB), Central Arnhem (CEA), Daly Basin (DAB), or Darwin Coastal (DAC) (Table S1), as defined by Interim Biogeographic Regionalisation for Australia (IBRA), version 7. Excised liver was stored in RNAlater™ Stabilization solution (Thermofisher Scientific, MA, USA), at room temperature while in the field and then at 4°C for longer term storage. Sampled animals displayed no obvious signs of serious pathology.

### 2.3 RNA extraction

Liver tissue was homogenized using the Qiagen Tissue Lyser II with 3mm stainless steel beads in Qiagen buffer RLT (Hilden, Germany); and RNA extracted using the Qiagen RNeasy Plus minikit (Hilden, Germany) according to the manufacturer’s protocol. Purified RNA was pooled in equimolar ratios into eleven pools, grouping RNA samples from the same lizard species and collection location, with six to twelve individuals per pool (Table 1, Table S1). Pooled RNA was further purified using the RNeasy MinElute clean-up kit (Qiagen, Hilden, Germany) and quantified using the Qubit RNA Broad-range Assay with the Qubit Fluorometer v3.0 (Thermofisher Scientific).

### 2.4 Meta-transcriptomic sequencing

RNA pools were assessed for quality using the Agilent 2100 Bioanalyzer with the Agilent RNA 6000 Nano Assay (Agilent Technologies, CA, USA). Library construction and sequencing was performed at the Australian Genomic Research Facility (Victoria, Australia). Libraries were constructed using the TruSeq Total RNA Library Preparation protocol (Illumina, CA, USA) following rRNA removal using the Illumina Ribo-zero Gold epidemiology kit. Paired-end (100 bp) sequencing of each RNA library was performed on a HiSeq 2500 sequencing platform (Illumina, CA, USA).

### 2.5 Genome/transcript assembly, annotation and abundance calculation

After trimming with Trimmomatic v0.38 (Bolger, Lohse and Usadel 2014), reads were assembled *de novo* using two separate assemblers – Trinity v2.5.1 (Grabherr et al. 2011) and Megahit v1.2.9 (Li et al. 2015) – to increase the chances of correctly assembling all viral contigs. Contigs from both methods were combined and duplicates removed (retaining the longest version) using CD-HIT-EST v4.8.1 (Fu et al. 2012). Viral contigs were identified using BLASTn (Altschul et al. 1990) and DIAMOND BLASTx (Buchfink, Reuter and Drost 2021) tools through alignment with the NCBI nucleotide (nt) database (*e*-value cut-off 1 x 10^-10^) and non-redundant protein (nr) database (*e*-value cut-off 1 x 10^-5^), respectively. The Geneious assembler (available in Geneious Prime ® 2022.2.2) was used to extend viral contigs where possible. Open reading frames were identified using the Find ORFs tool within Geneious Prime, and conserved domains were identified using RSP-TBLASTN v2.12.0+ (Altschul et al. 1997) against the NCBI Conserved Domains database. The abundance of each viral contig was calculated as expected counts (from mapped trimmed reads) using the RSEM tool in Trinity v2.5.2 (Li et al. 2010). Overall abundance was calculated as expected count/total number of trimmed reads in library x 100. Novel viruses that shared greater than 90% amino acid identity in the RdRp were considered to represent the same species. Likely vertebrate-infecting viruses were defined as those that belong to classically vertebrate-infecting viral families and/or those that clustered with viruses known to infect vertebrate species in phylogenetic trees.

### 2.6 Alpha and beta diversity analyses

Diversity statistics were only obtained for viruses considered likely to infect vertebrate species (i.e., those likely to infect the sampled host) and viruses likely to be exogenous. *Retroviruses* were excluded due to the difficulty in determining whether they are endogenous or exogenous, and viral groups with disrupted ORFs were considered likely to be endogenous. We performed generalized linear models and selected the best-fit model at both the infraorder and family taxonomic level (among a set of possible models describing the relationship between taxonomy, habitat, environment, range size and the number of individuals in the library) based on the lowest AIC, as done previously (Geoghegan et al. 2021; Chen et al. 2022). Host genus and species level were not considered due to small sample sizes. The Csex_M library was excluded from all analyses because it contained no exogenous, biologically relevant viruses and as the only riparian library, may have skewed results. All other libraries were included for comparisons considering host taxonomy at the level of infraorder. Omar_M was similarly removed for comparisons at the host family level as it was the only library representing Diplodactylidae. Alpha diversity, including richness, Shannon index, Simpson Index, Shannon effective and Simpson effective were calculated for each library (Lagkouvardos et al. 2017). Beta diversity was calculated using the Bray–Curtis dissimilarity matrix and virome structure was plotted as a function of nonmetric multidimensional scaling (NMDS) ordination and tested using Adonis tests (PERMANOVA) using the *vegan* and *phyloseq* packages (McMurdie and Holmes 2013; Oksanen et al. 2022). Analyses were performed in R version 4.2.3 in R Studio 2022.07.1.

### 2.7 Phylogenetic analysis

Viral amino acid sequences were aligned with representative sequences from the same viral family obtained from NCBI, using Clustal Omega v1.2.3 available in Geneious Prime. Where necessary, large data sets were condensed to a more manageable size using CD-HIT version 4.8.1 (Fu et al. 2012). TrimAl v1.4.1 (Capella-Gutiérrez, Silla-Martínez and Gabaldón 2009) was used to remove ambiguously aligned regions (using the gappyout setting in all cases except for the *Flaviviridae* alignment which required stricter manually applied settings to remove uninformative columns). Alignments were visualized in Geneious Prime. Maximum likelihood trees were inferred using IQTree v2.1.3 (Nguyen et al. 2015) with model selection estimated using ModelFinder within IQTree (Kalyaanamoorthy et al. 2017). Branch supports were estimated with the Shimodaira-Hasegawa (SH)-like approximate likelihood ratio test (Guindon et al. 2010). The size and length of each alignment is provided in Table S2 and details of assembled viral nucleotide consensus sequences that were translated for inclusion in phylogenies are provided in Table S3.

To reduce the reporting of false positives due to index-hopping during sequencing for each viral contig present in more than one library, a viral contig was presumed to be a contaminant from another library if it met the following criteria: contig abundance was less than 0.1% of the abundance of that contig in the library where that contig was most abundant. This is based on the index-hopping rate of about 0.1-2% as listed by Illumina (https://sapac.illumina.com/techniques/sequencing/ngs-library-prep/multiplexing/index-hopping.html).

### 2.8 Virus nomenclature

Novel viral species were determined based on demarcation criteria assigned by the ICTV (International Committee on Taxonomy of Viruses) for the relevant families/genera (https://ictv.global/report/genome) and were randomly assigned provisional names. Note that because these virus names are provisional, we have only assigned common names rather than a full binomial nomenclature.

## 3. Results

### 3.1 Viruses in Australian lizard species

We sequenced 11 pooled RNA libraries with an average read count of 35,041,464 (Table 1). Each library was generated from liver samples of 6 to 12 apparently healthy lizards of a single species sampled from either the mesic Top End or from the relatively arid east Kimberley regions of Australia (Victoria-Bonaparte region; Table 1). Two widespread gecko species – *Heteronotia binoei* and *Gehyra nana* – were sampled from both regions. A high diversity of vertebrate-associated viruses was detected, comprising nine different families and 12 genera (Figure 1). While for most viral families only a single genus and species were documented, we identified viruses from at least two genera in each of the *Flaviviridae*, *Picornaviridae*, *Rhabdoviridae* and *Amnoonviridae,* and two species of the same genera in the *Astroviridae* (Figure 1B). Despite being present in only one library, the *Arenaviridae* were by far the most abundant viral family (Figure 1), with the *Flaviviridae* the second most abundant and detected in all three lizard families (5/11 libraries) (Figure 1A). Members of the *Picornaviridae* were also abundant and detected in five of 11 libraries, although were restricted to the Gekkota species (Gekkonidae and Diplodactylidae). No vertebrate-infecting viruses were shared between different lizard species (Figure 1B). At higher taxonomic scales, the genera *Hepacivirus* (*Flaviviridae*), *Shanbavirus* (*Picornaviridae),* and an unclassified genus of *Astroviridae*, were found in multiple lizard species (Figure 1). Five viral families were found across multiple host species: the *Flaviviridae*, *Picornaviridae*, *Astroviridae*, *Rhabdoviridae* and *Amnoonviridae*.

**Table 1.**
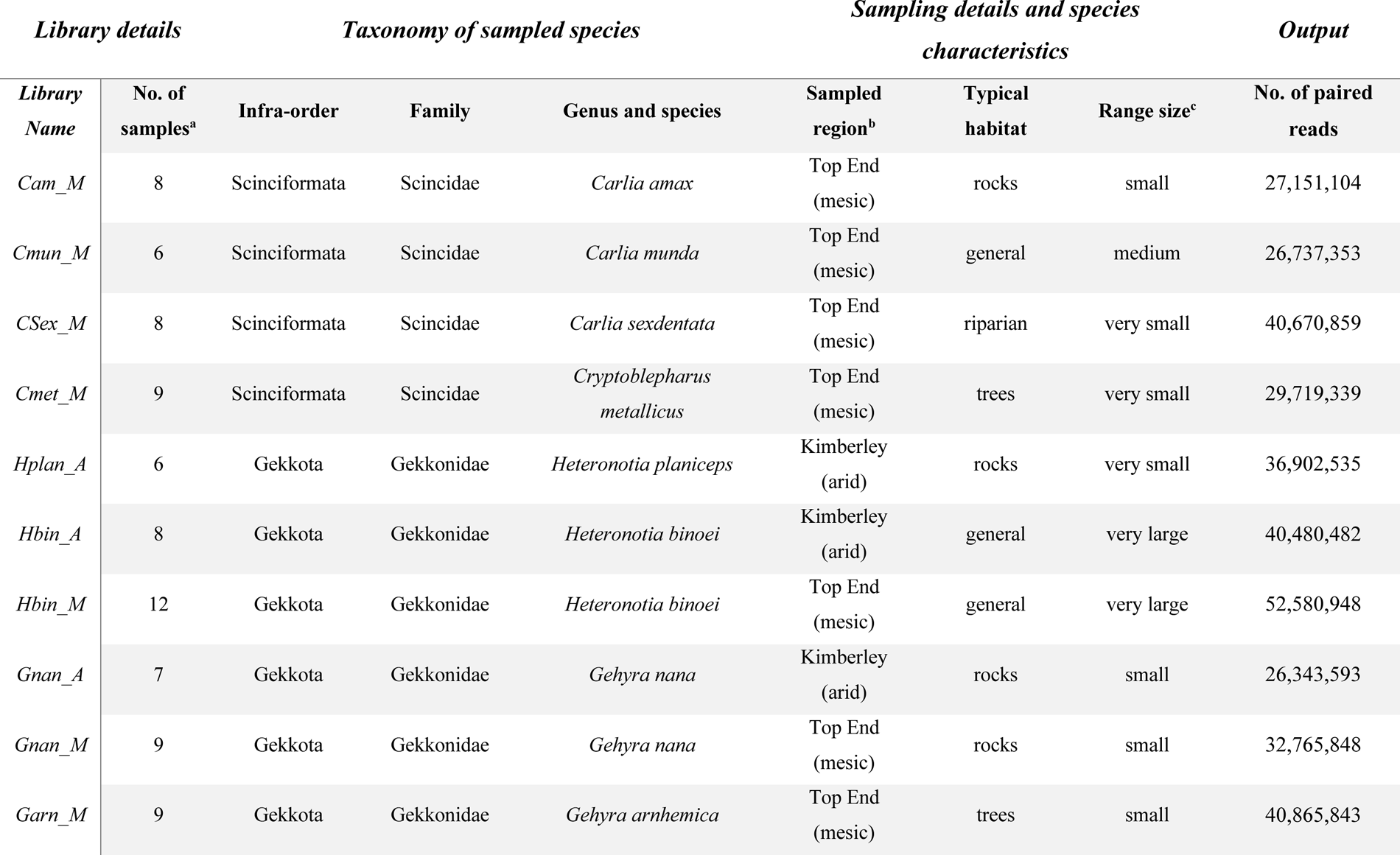
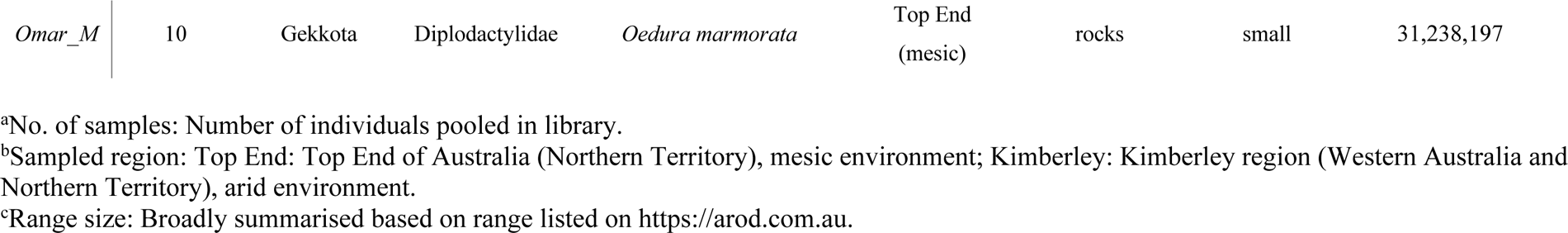
Details of lizard livers sampled and sequencing output for each library (see Table S1 for additional details).

**Figure 1.**
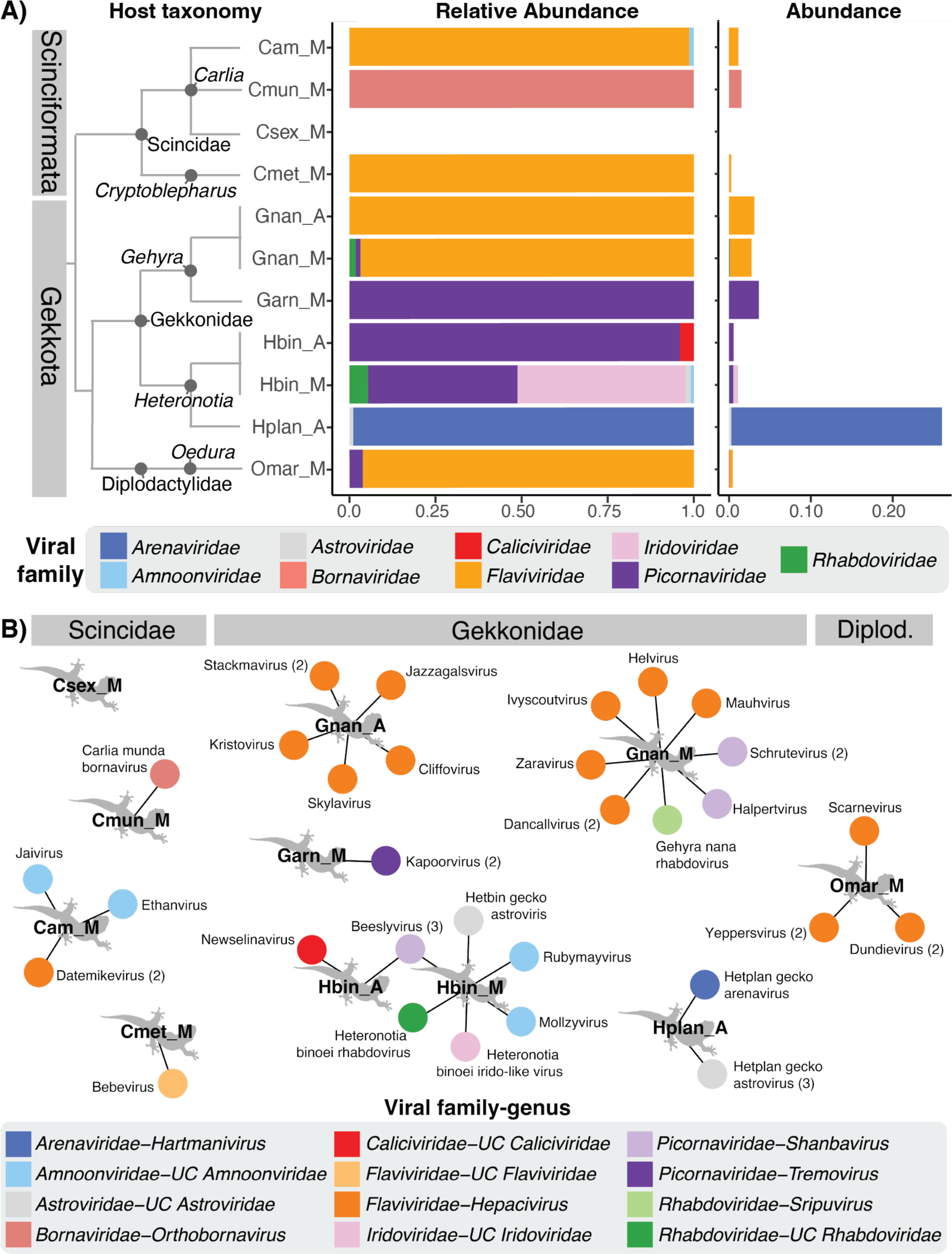
Biologically relevant viruses (i.e., viruses that infect vertebrates) in the sampled lizard species. (A) Relative abundance (left plot) and overall abundance (right) of the vertebrate-infecting virus families present in each library. Libraries are plotted in taxonomic sequence, with host relationships indicated by a cladogram and host infraorder indicated in grey bars. Library names are as follows: Cam_M, *Carlia amax (collected from a mesic environment)*; Cmun_M, *Carlia munda* (mesic), Csex_M, *Carlia sexdentata* (mesic); Cmet_M, *Cryptoblepharus metallicus* (mesic); Gnan_A, *Gehyra nana* (arid); Gnan_M *Gehyra nana* (mesic); Garn_M, *Gehyra arnhemica* (mesic); Hbin_A, *Heteronotia binoei* (arid); Hbin_M, *Heteronotia binoei* (mesic); Hplan_A, *Heteronotia planiceps* (arid); Omar_M, *Oedura marmorata* (mesic). (B) Viruses found in each library are represented by circles coloured by viral genus (UC=unclassified genus), with lines connecting them to the libraries in which they were found. Libraries are grouped by host family (indicated by grey bars above; Diplod.=Diplodactylidae). Numbers in parentheses beside virus names indicates the number of variants of that virus detected (where >1). Note that all the *Amnoonviridae* belong to currently unclassified genera, but may represent more than one genus.

A number of other vertebrate-associated viral families were detected, but were determined as likely endogenous virus elements (EVEs) because longer contigs had disrupted open reading frames. These included members of the *Adintoviridae*, *Hepadnaviridae*, *Retroviridae*, and *Circoviridae,* and were excluded from abundance and alpha diversity analyses. Of note, library CSex_M (comprising *Carlia sexdentata*) did not contain any biologically relevant viruses, aside from some likely EVEs, but did contain some viruses that are not associated with vertebrates (Figure S1). Across all libraries, we detected several viral groups that were unlikely to be associated with vertebrate infection, representing the *Permutotetraviridae*, *Chuviridae*, *Tectiviridae*, *Mimiviridae*, *Autolykiviridae*, *Baculoviridae*, unclassified Ortervirales, unclassified Riboviria, and tombus-like, solemo-like, narna-like, partiti-like, and toti-like viruses. None of these viruses were abundant (Figure S1) and were considered likely to be viruses of commensal organisms in the lizard livers (i.e., narna-like) or contaminants. Only vertebrate-associated viral families are included in the analyses described below.

### 3.2 Abundance and diversity of lizard viruses

To help determine the factors influencing virome diversity, we compared the abundance and alpha diversity in biologically relevant viruses between libraries from different lizard species and sampling regions. Specifically, we considered host taxonomy (at the level of infraorder and family) as well as a range of ecological variables, including habitat, environment and range size (Csex_M was excluded from these analyses as it was the only riparian library and no biologically relevant viruses were found in this library). Given variability in the number of individuals per pool, we first assessed the effect of the number of samples per pool/library to account for any bias introduced by sampling strategy. The number of samples per pool correlated with the Richness (p=0.037) and Shannon diversity (p=0.015), but not the abundance (p= 0.140) of vertebrate viruses, or the Shannon Effective (p=0.052), Simpson (p=0.434) or Simpson Effective diversity (p=0.077) (Figure S2). As such, the number of samples per pool was considered in all statistical models, and included as cofactors in final models addressing Richness and Shannon diversity.

Host taxonomy and habitat were consistently important parameters in the best-fit models for all diversity measures regardless of the level of taxonomy considered, indicating that these factors are important modulators of viral diversity in this sample. Taxonomy had a statistically significant effect on viral richness regardless of taxonomic level (host infraorder p=0.030, family p=0.002; Figure 2A and 2C), but was not significant in modulating abundance or Shannon and Simpson indexes (Figure S3). Habitat significantly modulated richness and Shannon index diversity when considering host infraorder (p=0.049 and p=0.025, respectively) and host family (p=0.002 and p=0.048; Figure 2B and 2D), and also had a significant effect on the Simpson index of diversity, but only when considering host infraorder (p=0.037; Figure S4). Range size was significant in the best-fit model for richness, regardless of the taxonomy level explored (host infraorder p=0.049, host family p=0.002).

**Figure 2.**
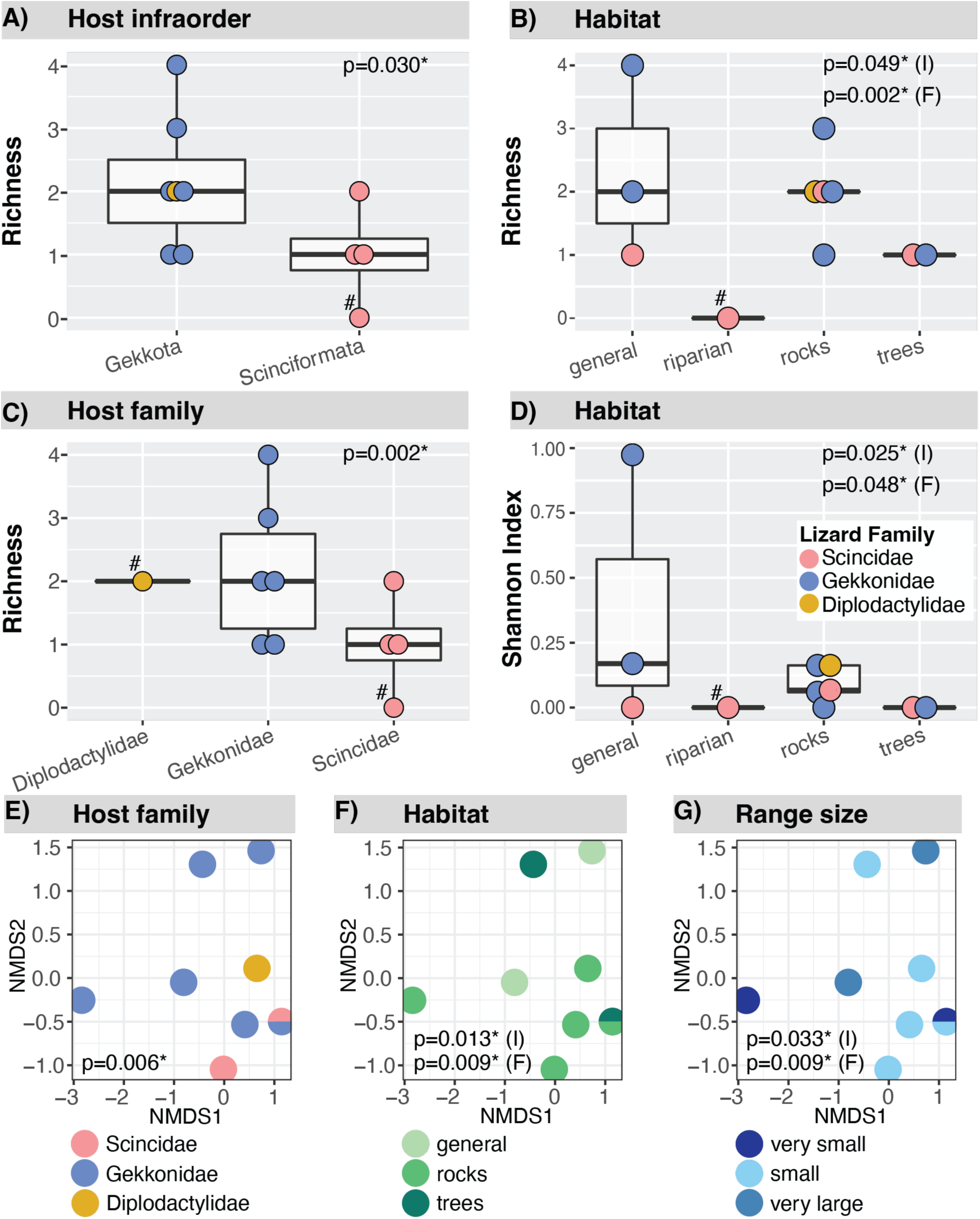
Diversity of biologically relevant viruses by host taxonomic and ecological variables. (A—D) viral alpha diversity plots: (A) viral richness according to host infraorder; (B) viral richness according to lizard habitat type; (C) viral richness according to host family; (D) Shannon index according to host habitat. Points are coloured by host family. Host taxonomic levels comprising only a single library were included in plots for visualisation only and excluded from the final statistical models (denoted by a # in host taxonomy alpha diversity plots). Csex_M was not included in statistical models or beta diversity plots as no biologically relevant viruses were found in this library and it was the only riparian library. (E—G) viral beta diversity NMDS plot coloured by: (E) host family, (F) host habitat, and (G) host range size. Cmet_M and Gnan_A overlap, depicted by a half circle. In all plots, P-values for habitat or range size when considered with taxonomy at the level of host infraorder are indicated with an “I”, while those considered with taxonomy at the level of host family are indicated with an “F”. P-values <0.05 were considered significant and indicated with an asterisk.

We also explored beta-diversity in relation to host traits (Figure 2E-G and Figure S5). A similar trend to alpha diversity was observed, in which taxonomy, habitat and range size were statistically significant in models considering host family (host family p=0.006, habitat p=0.009, range size p=0.009), as was habitat and range size in models considering host infraorder (p=0.013 and 0.033, respectively). Overall, we documented a general trend of host taxonomy, habitat and range size as potential modulators of viral diversity.

### 3.3 Evolutionary relationships of novel viruses

#### 3.3.1 Flaviviridae

Strikingly, most of the *Flaviviridae* identified in the lizards studied here fell within the genus *Hepacivirus*. The Australian lizard hepaciviruses were all novel and fell into three clades in a host specific manner (Figure 4). Hepaciviruses from *Carlia amax* formed a monophyletic group with a viviparous lizard hepacivirus from *Zootoca vivipa*ra (Lacertidae) sampled in the UK (Scotland), and a Gecko hepacivirus previously sampled from *Oedura* in Australia (although the RdRp sequence from the Gecko hepacivirus was only 92 amino acid residues, necessarily impacting phylogenetic accuracy). In addition, the long branch lengths suggest substantial unsampled genetic diversity within this clade. While this clade has strong support, its specific phylogenetic placement within the hepaciviruses is poorly resolved such that its closest relatives are unclear (Figure 4). *Gehyra nana* had a high diversity of hepaciviruses, with viral sequences clustering into two groups: one distinct lineage that broadly clustered with mammalian viruses and a second that grouped with reptile and bird viruses (Figure 4). Viruses in the avian/reptile lineage grouped with viruses from other Gekkota species sampled in China, including a *Goniurosaurus luii* (Gekkota; Eublepharidae, AVM87252.1), *Hemidactylus bowringii* (Gekkota; Eublepharidae, AVM87554.1) and a *Teratoscincus roborowskii* (Gekkota; Sphaerodactylidae, AVM87251.1). This is consistent with the Asian origins of *Gehyra* (Oliver and Hugall 2017). Viruses from the *Oedura marmorata* sampled in this study formed their own clade within the avian/reptile host lineage, basal to those from China and *G. nana*.

Interestingly, the *Flaviviridae* identified in the *Cryptoblepharus metallicus* library did not cluster with the other Australian lizard *Flaviviridae*, but rather with Tanama bat virus and Lumpfish flavivirus (Figure 4), which are part of a broad “flavi-like” group that are phylogenetically distinct from members of the genus *Orthoflavivirus*. The long branch lengths between these three viruses suggest that there is considerable unsampled viral diversity in this clade, and that this constitutes a new virus that we have tentatively named Bebevirus.

**Figure 3.**
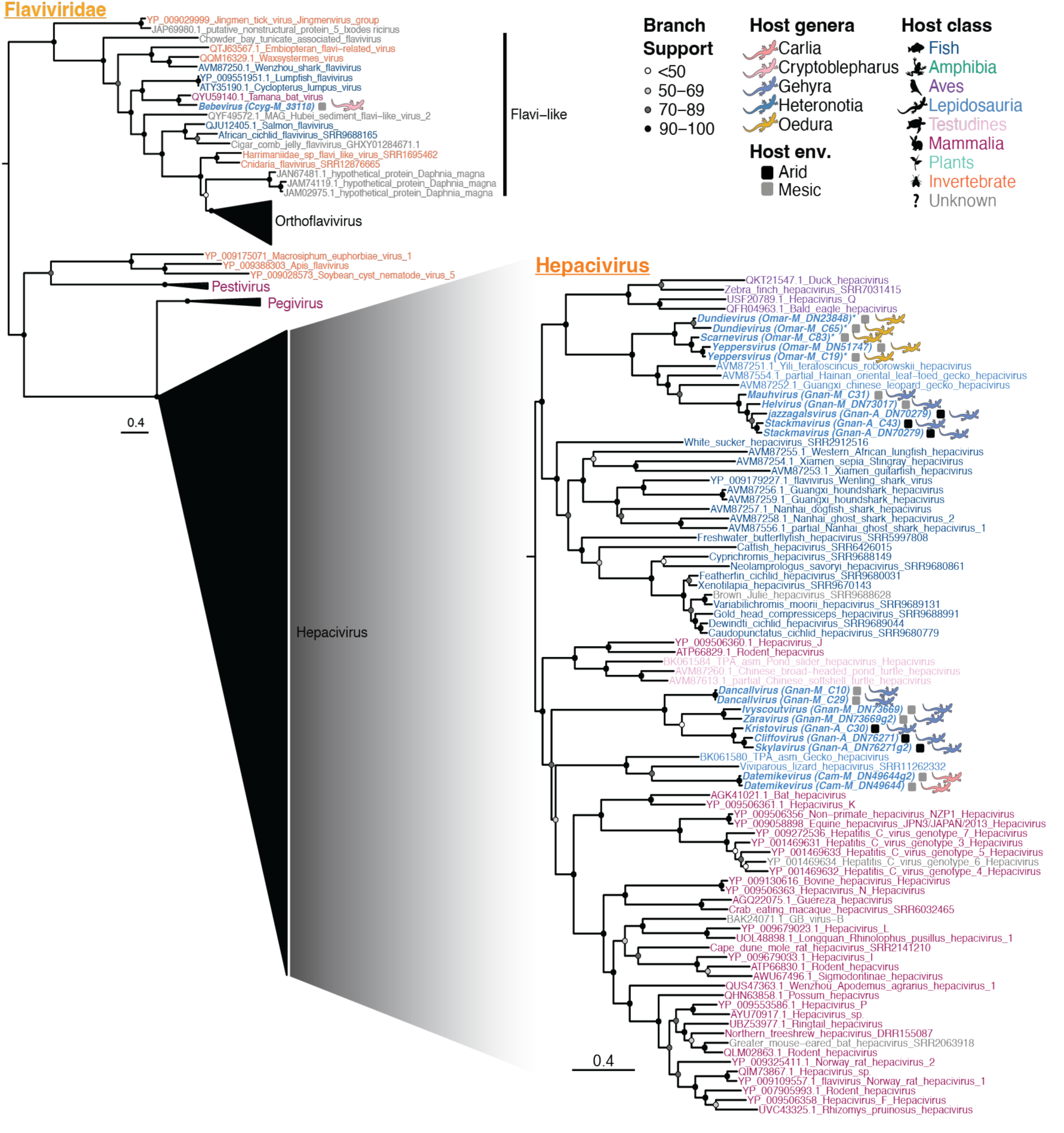
Maximum likelihood phylogeny of the RdRp of *Flaviviridae* in Australian lizards. Taxon names are coloured according to apparent host. Viruses discovered in this study are indicated by bold and italicized taxa names and lizard silhouettes beside taxa names, coloured by host genus. Squares next to the taxa names indicate the sampling location/environment (env.) for viruses discovered in this study. An asterisk beside the taxa name for viruses detected here indicates that the sequence is not the complete length of the alignment. The accession number is indicated in the taxon name for sequences downloaded from NCBI. Circles at the nodes represent the branch support as estimated using the SH-like approximate likelihood ratio test. Trees are mid-point rooted for clarity. Scale bars indicate the number of amino acid substitutions per site. To achieve greater resolution, the *Hepacivirus* phylogeny was estimated from a sequence alignment of this genus only.

#### 3.3.2 Picornaviridae

Members of the *Picornaviridae* were found in all Gekkota species analyzed, with the exception of *H. planiceps* (although a picornavirus contig from *O. marmorata* was excluded from the phylogeny as there were no RdRp contigs of sufficient length). The picornaviruses isolated from lizards in this study fell into two main groups. Viruses from *H. binoei* and *G. nana* grouped together and shared a common ancestor with bat picornavirus (AIF74248.1; 53.5-62% amino acid [aa] identity) and Chameleon picornavirus 1 (DAZ91100; 50-58.7% aa identity) (Figure 4), a virus found in multiple lizard species including *Kinyongia boehmei* (Chamaeleonidae), *Podarcis muralis* (Lacertidae), and *Timon pater* (Lacertidae). Bat picornavirus is a member of the genus *Shanbavirus*, and (based on identity in the RdRp) the *H. binoei* and *G. nana* viruses, as well as Chameleon picornavirus 1, should also belong to the shanbaviruses (https://ictv.global/report/chapter/picornaviridae/picornaviridae). Three distinct *Shanbavirus* species were identified in this study, provisionally named Halpertvirus (*G. nana* host), Schrutevirus (*G. nana* host), and Beeslyvirus (*H. binoei* host). Notably, Beeslyvirus was found in *H. binoei* from both mesic and arid environments and was the only virus found in more than one library.

The second group contained the *G. arnhemica* picornavirus sequences, which were closely related and hence assigned as the same virus species, here provisionally named Kapoorvirus. This virus was most closely related to worm lizard picornavirus (57.6% aa identity in the RdRp) from *Blanus cinereus* (Blanidae) and more broadly related to members of the genus *Tremovirus* (Figure 4), associated with reptiles and birds, and known to cause encephalomyelitis in avian hosts. The Kapoorviruses shared 34.3-41.6% amino acid identity in the RdRp with the *Tremoviruses* included in the phylogeny, which placed them on the border of the level of divergence required for a new genus (64% divergence). As such, we have tentatively placed them within the genus *Tremovirus*.

**Figure 4.**
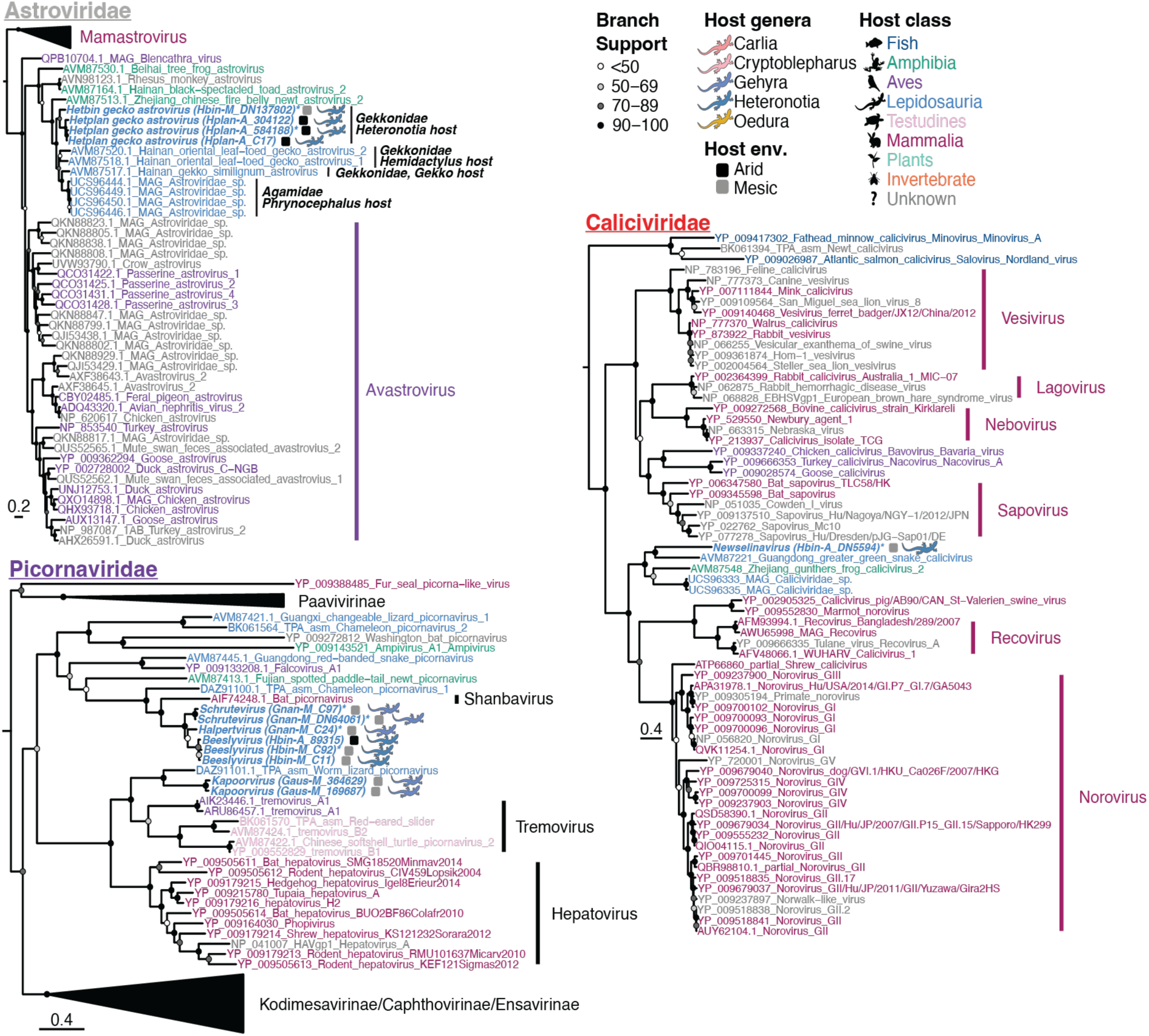
Maximum likelihood phylogenies of the RdRp of positive-sense RNA viruses in Australian lizards (with the exception of the *Flaviviridae*, shown in Figure 3). Taxon names are coloured according to apparent host. Viruses discovered in this study are indicated by bold and italicized taxa names and lizard silhouettes beside taxa names, which are coloured by host genus. Squares next to the taxa names indicate the sampling location/environment (env.) for viruses discovered in this study. An asterisk beside the taxa name for viruses detected here indicates that the sequence is not the complete length of the alignment. The accession number is indicated in the taxon name for sequences downloaded from NCBI. Circles at the nodes represent the branch support as estimated using the SH-like approximate likelihood ratio test. Trees are mid-point rooted for clarity. Scale bars indicate the number of amino acid substitutions per site. Host family and genus names are indicated for the Lepidosauria in the *Astroviridae* phylogeny to demonstrate virus-host co-divergence within the Lepidosauria.

#### 3.3.3 Astroviridae

Members of the *Astroviridae* were detected in two *Heteronotia* libraries from two different species – *H. binoei* and *H. planiceps.* A distinct astrovirus was identified in each library, designated as Hetplan gecko astrovirus (from *H. planiceps*) and Hetbin gecko astrovirus (from *H.* binoei), and the two viruses grouped together in the RdRp phylogeny (Figure 4). These viruses then grouped more broadly with a larger well-supported clade of viruses from lizards (Figure 4), clustering in a fashion reflective of host family and genus. Specifically, the viruses from *Heteronotia* grouped closely with viruses from other members of the Gekkonidae, including *Hemidactylus bowringii* and *Gekko similignum,* with each also clustering by host genus, all of which fell basal to a clade of viruses from Agamidae (*Phrynocephalus erythrurus* and *Phrynocephalus theobaldi*), mirroring the host phylogeny (Figure 4). Genus demarcation criteria for the *Astroviridae* is unclear and is most strongly associated with host species. As a consequence, the viruses found here, along with the published lizard astroviruses, would most likely form their own genus. It is also notable that the lizard astrovirus cluster grouped more broadly with viruses found in amphibians and birds.

#### 3.3.4 Rhabdoviridae

Viral contigs related to the *Rhabdoviridae* were identified in two libraries (Figure 1). The *H. binoei* rhabdovirus fell with a group of viruses not classified to any genus, including those isolated from ray-finned fishes (*Actinopterygii*; from liver, gut and gills), and a spotted paddle-tail newt (*Pachytriton brevipes*; from gut) (Figure 5). This virus is highly divergent (49% aa identity in the RdRp to the closest relative, Wenling dimarhabdovirus 9) and would therefore represent a new species, that we have named Heteronotia binoei rhabdovirus. This virus, together with Wenling dimarhabdovirus 9, Fujian dimarhabdovirus, and Wenling dimarhabdovirus 1, should form at least one new genus based on the observed level of sequence diversity. This group of unclassified viruses then fell as a sister-group to a large clade of rhabdoviruses including lyssavirus and many other pathogenic viruses (Figure 5).

Only short *Rhabdoviridae* contigs could be obtained for the RdRp region from the *G. nana* library (Gnan_M), the longest being 124 residues. A phylogeny was estimated using a smaller region of the RdRp (230 aa after trimAl trimming) where this contig aligned to confirm the phylogenic groupings observed in the larger alignment (774 aa after trimAl trimming). Regardless of the RdRp alignment used, the *G. nana* rhabdovirus grouped with the *Sripuviruses* (Figure 5). It shared 75% identity in the RdRp protein to its closest relative – Almpiwar virus, isolated from skinks in Northern Queensland Australia. As all RdRp contigs from Gnan_M (including the one in the phylogeny) exhibited less than 90% amino acid identity to the closest relative, this virus likely represents a new species of *Sripuvirus*, tentatively named Gehyra nana rhabdovirus.

**Figure 5.**
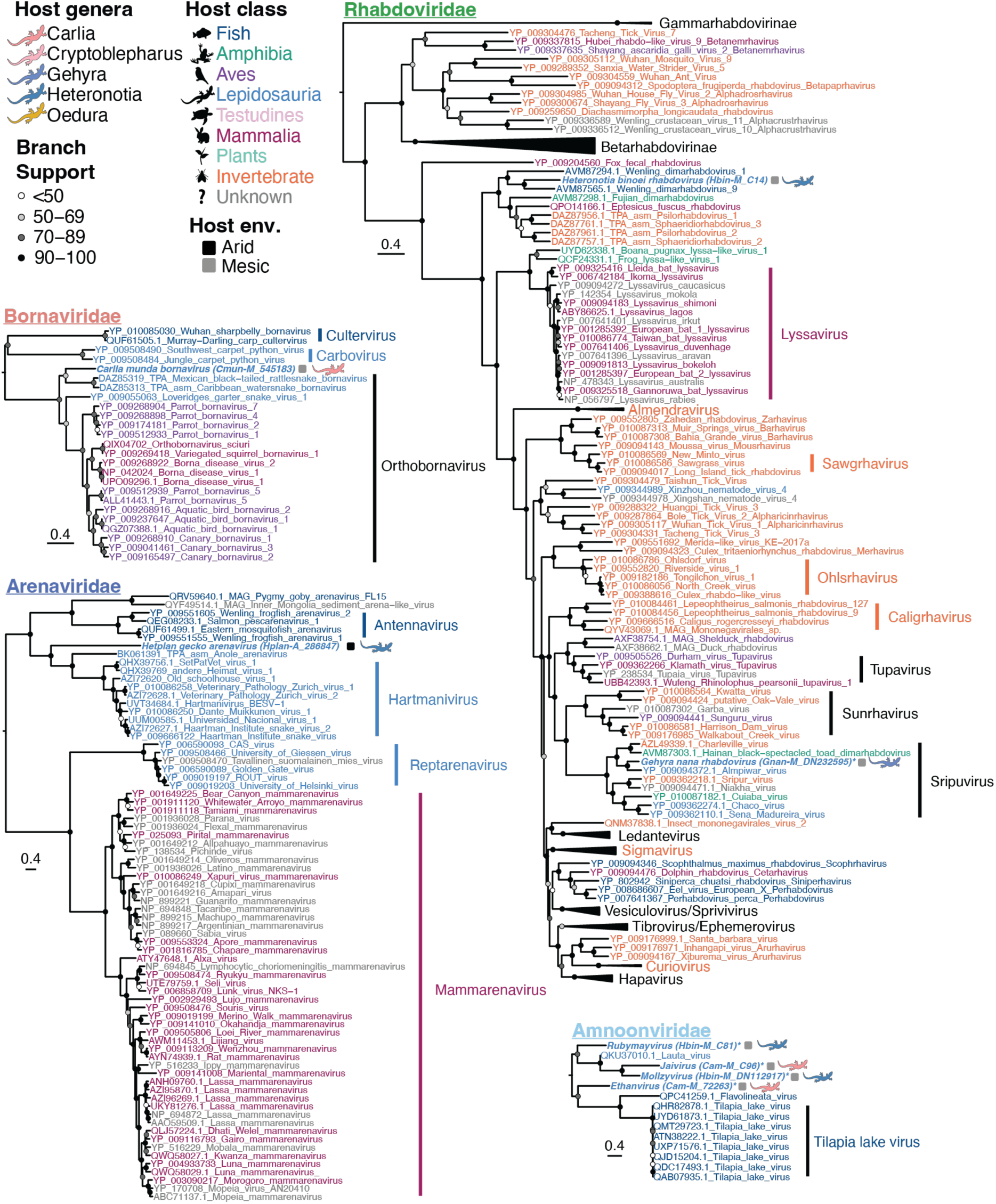
Maximum likelihood phylogenies of the RdRp of negative-sense RNA viruses in Australian lizards. Taxon names are coloured according to apparent host. Viruses discovered in this study are indicated by bold and italicized taxa names and lizard silhouettes beside taxa names, which are coloured by host genus. Squares next to the taxa names indicate the sampling location/environment (env.) for viruses discovered in this study. An asterisk beside the taxa name for viruses detected here indicates that the sequence is not the complete length of the alignment. Accession numbers are indicated in the taxon name for sequences downloaded from NCBI. Circles at the nodes represent the branch support as estimated using the SH-like approximate likelihood ratio test. Trees are mid-point rooted for clarity. Scale bars indicate the number of amino acid substitutions per site.

#### 3.3.5 Amnoonviridae

Several contigs related to Lauta virus (MT386081), a member of the *Amnoonviridae*, were identified in *C. amax* and *H. binoei* sampled from the same area of the north-east Top End. However, the contigs obtained were short and hence difficult to classify. The amino acid sequence identity between the longest contigs (270 aa and 238 aa) from the two libraries is only 26.7%, while the identity of the longest contigs from *C. amax* and *H. binoei* with the nearest published relative (MT386081, Lauta virus) were 31.3% and 35.8%, respectively. The nearest known relative, found in an Australian gecko (*Gehyra lauta*) liver in 2013 (Ortiz-Baez et al. 2020), is itself highly divergent from other published viruses, and was only detected using a protein structure prediction approach. All *Amnoonviridae* contigs detected in this study were from the RdRp segment as these viruses are so distinct from known viruses that it was challenging to identify other genomic regions, particularly as these viruses are likely to be segmented. Phylogenetic analysis of the RdRp of contigs longer than 100 aa from each library suggested that there were four distinct species of lizard virus within *Amnoonviridae*, two in each library (Figure 5). Three of these species clustered with Lauta virus, creating a lizard-infecting clade, while the fourth was basal (yet quite distant) to a clade of fish viruses, including Tilapia lake virus (Figure 5). However, the clustering of these contigs may be unreliable since they are short (160 – 270 aa) and not the complete length of the RdRp. Additionally, contigs from the same library did not overlap (or only overlapped minimally) in the alignment and therefore their relationship is difficult to determine.

### 3.4 Viral families in which a single species was identified

A single virus species was detected in the lizard metatranscriptomes from the *Caliciviridae, Arenaviridae*, *Bornaviridae*, and *Iridoviridae*. A member of the *Caliciviridae* was detected in the *H. binoei* (Hbin_A) (Figure 1), but only partial contigs were obtained which aligned to most of the capsid, parts of the RdRp, 2C-like protein and the proteinase. Alignment of a 449 aa region of the RdRp with published sequences grouped this virus with caliciviruses from other squamates; *Cyclophiops major* (Colubridae), *Phrynocephalus theobaldi* (Agamidae), and a frog, *Sylvirana guentheri* (Ranidae) (Figure 4). The *H. binoei* calicivirus was highly divergent from its closest relative – Guangdong greater green snake calicivirus (AVM87221) – sharing only 35.4% identity in the aligned region of the RdRp. We therefore suggest that it comprises a new species that we have tentatively named Newselinavirus. Based on alignment of the near complete major capsid protein (527 aa), this virus has >60% amino acid sequence difference to other caliciviruses and therefore would also constitute a different genus (https://ictv.global/report/chapter/caliciviridae/caliciviridae).

A member of the *Arenaviridae* was detected in *H. planiceps* (Hplan_A) with the highest abundance of any virus detected in this study (>0.2% of total reads for that library, Figure 1) with a near complete genome obtained. This virus clusters with the *Hartmaniviruses* (Figure 5), a genus known to infect snakes with unknown pathogenicity, although with only 23.6% aa identity to its closest relative in the RdRp region. However, based on the genus demarcation criteria for *Arenaviridae* (i.e., members of the same genus share >35% nucleotide identity in the L gene; https://ictv.global/report/chapter/arenaviridae/arenaviridae), this virus falls just within the genus *Hartmanivirus*. Like other members of this genus, the virus detected here, tentatively named Hetplan gecko arenavirus, lacked the gene encoding the zinc-binding matrix protein (Hepojoki et al. 2015).

A near-complete genome of a member of the *Bornaviridae* was found in the *C. munda* library (Cmun_M). Phylogenetic analysis of the RdRp showed that it formed a clade with two snake viruses from the genus *Orthobornavirus* which also contained Loveridges garter snake virus 1 (YP_009055063) and viruses with mammalian and avian hosts (Figure 5). As it shared only 53.5% nucleotide identity across the entire genome with its nearest relative, the virus detected here would constitute its own species (Kuhn et al. 2015), tentatively named Carlia munda bornavirus, and would likely belong to the genus *Orthobornavirus*. It is not clear whether the two snake viruses that grouped with Carlia munda bornavirus are pathogenic (Pfaff and Rubbenstroth 2021), but other members of the *Orthobornavirus* clade cause neurotropic disease in mammals and avian hosts, while members of the *Carbovirus* genus cause neurological disease in snakes (Hyndman et al. 2018).

Transcripts from multiple genes related to the *Iridoviridae* (double-strand DNA viruses) were detected in *H. binoei* (Hbin_M) (Figure 1), including the major capsid, myristylated membrane protein, phosphotransferase, ATPase, DNA helicase, dUTP, and NTPase I. This suggested that an *Iridoviridae* was actively replicating in one or more lizards sampled in this library. Phylogenetic analysis of the major capsid protein revealed that this virus clustered within the subfamily *Betairidovirinae* that was originally considered invertebrate-specific but which is increasingly being associated with vertebrate infection (Russo et al. 2021) (Figure 6). Interestingly, this virus grouped with erythrocytic necrosis virus, a pathogen of fish (72.4% aa identity in the capsid), and Rhinella marina erythrocytic-like virus found in cane toads (85.4% aa identity) and would likely fall into the same genus (https://ictv.global/report/chapter/iridoviridae/iridoviridae), but as a novel species that we have named Heteronotia binoei irido-like virus (Figure 6). A new genus containing these and erythrocytic viruses of ectothermic hosts has been putatively proposed, but not yet formally ratified or named (Emmenegger et al. 2014). Three other erythrocytic viruses from Squamata hosts – Thamnophis sauritus erythrocytic virus (EV), Lacerta monticola EV, and Pogona vitticeps EV – fell within this erythrocytic virus lineage based on sequence of a short PCR product of a homolog to a region of the DNA dependent DNA polymerase (Grosset et al. 2014; Russo et al. 2018), although the transcript encoding this particular protein was not found in our data, precluding a phylogenetic analysis. However, it is likely the virus discovered here is a relative to the other Squamata erythrocytic viruses and supports the presence of Squamata hosts for this lineage.

**Figure 6.**
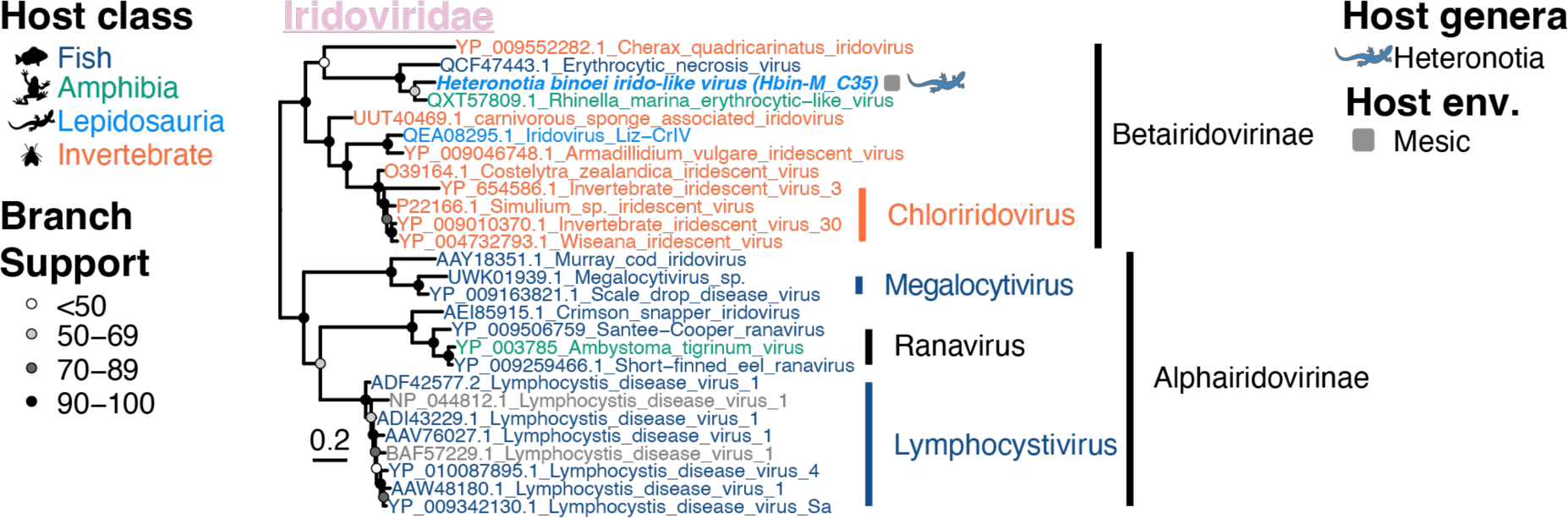
Maximum likelihood phylogenetic tree of the major capsid protein of the *Iridoviridae* in Australian lizards. Taxon names are coloured according to apparent host. The virus discovered in this study is indicated by a bold and italicized taxon name and a lizard silhouette beside the taxon name, which is coloured by host genus. The square next to the taxon name indicates the sampling location/environment (env.) for the virus discovered in this study. The accession number is indicated in the taxon name for sequences downloaded from NCBI. Circles at the nodes represent the branch support as estimated using the SH-like approximate likelihood ratio test. The tree is mid-point rooted for clarity, and the scale bar indicates the number of amino acid substitutions per site.

## 4. Discussion

We used Australian lizards as model organisms to better understand the host determinants of viral abundance and diversity, while simultaneously expanding our knowledge of the lizard virome and virus evolution. Indeed, to date relatively little attention has been paid to the viromes of squamates (lizards and snakes) (Shi et al. 2018; Harding et al. 2022). In Australia, the Squamata are the most species-rich vertebrate assemblage (Wilson and Swan 2017) and the varied ages of Squamate radiations (Brennan and Oliver 2017), alongside their varied population structures and their adaption to multiple habitats and environments (Pianka 1969; Pianka 1973), makes them an informative group to study in the context of viral evolution and ecology. Using a meta-transcriptomics approach we characterized the virome of nine lizard species from three families and five genera, finding a wide diversity of novel viruses, including those from families/genera that commonly cause disease in humans and other animals, including the *Flaviviridae* (*Hepacivirus*), *Bornaviridae*, *Rhabdoviridae*, and *Picornaviridae*. Our findings expand many known viral families, adding entire new clades. This includes a new lizard-specific clade of the *Amnoonviridae*, a family previously containing only fish viruses and a single lizard-infecting virus (Ortiz-Baez et al. 2020).

Viral diversity and abundance differed markedly between hosts, and consistent with other studies on virome and pathogen ecology, environmental variables had some effect on virome diversity across our sampled hosts – particularly host habitat. The effect of various habitats on alpha diversity in viruses has previously been demonstrated in fish (Geoghegan et al. 2021), rodents (Tirera et al. 2021), and bats (Bergner et al. 2020), while virome composition has also been shown to differ significantly in different habitats for bats, shrews and rats (Chen et al. 2023). As different habitats have a range of environmental factors that modulate host behavior and ecology, it is unsurprising that host habitat may also impact virome composition. For example, foraging behavior is likely to play a role in the transmission of low pathogenic avian influenza, where avian species that forage in shallow water are more likely to influence avian influenza ecology (Wille et al. 2023).

Host taxonomy also impacted measures of virus richness and beta diversity, consistent with some degree of host-specific virus evolution, including co-divergence. The effect of host taxonomy on viral diversity is similar to that seen in fish (Geoghegan et al. 2021) and mammalian viromes (Olival et al. 2017), although a different pattern was observed in some bird viromes (Wille et al. 2018, 2019), suggesting that it is dependent on the taxa in question. Over evolutionary time the likelihood of successful cross-species virus transmission is predicted to decline because of host genetic differences in viral receptors, the cellular machinery required for replication, immune response and other factors that affect viral infection and spread (Ferris, Heise and Baric 2016). This may in part explain the link between host taxonomy and viral richness and composition. Alternatively, this association could simply reflect virome-level virus-host co-divergence over longer evolutionary timescales. Indeed, in most cases, the lizard viruses discovered here grouped together and/or with viruses found in other members of the reptilian class Lepidosauria, and phylogenies generally followed a broad pattern of host-virus co-divergence. For example, in the case of the *Astroviridae*, virus sequences from Squamata clearly group by host genera, and viruses from the Gekkonidae fall basal to those from the Agamidae, matching the host phylogeny (Oliver and Hugall 2017). Similarly, in the case of the relatively well-sampled hepaciviruses and picornaviruses, more closely related Squamata hosts tended to carry more closely related viruses, with viruses from the same host species forming a monophyletic group and viruses from the same family and infraorder similarly tending to do the same.

As well as providing some evidence for virus-host co-divergence, we documented many instances of virus-host jumping or multi-host viruses at higher taxonomic scales. Indeed, in some cases, lizard viruses clustered more closely with amphibian, mammalian or fish viruses, rather than with other reptile or bird viruses. Three distinct clades of lizard hepaciviruses were identified, all of which group separately to a clade of turtle hepaciviruses. Although the hepacivirus phylogeny is difficult to resolve, the presence of four clades of reptile hepaciviruses and the placement of two reptile clades within the mammalian hepacivirus clade is suggestive of host jumping between reptiles and mammals at some time in the distant evolutionary past. Similarly, the *Iridoviridae* and *Rhabdoviridae* phylogenies are highly incongruent with their host phylogenies, suggesting frequent host jumping. In the case of the two lizard *Rhabdoviridae*, one groups with fish viruses while the other clusters in the *Sripuvirus* clade with viruses from reptiles, amphibians and invertebrates. The closest relative of the *Sripuvirus* detected here, Almpiwar virus, was isolated from skinks in Northern Queensland Australia, and neutralizing antibodies against Almpiwar virus were also found in the sera of crocodiles, a wild bird and multiple mammalian hosts, including a human (McAllister et al. 2014). Although, the presence of neutralizing antibodies against Almpiwar virus in mammals was relatively rare, it does suggest that this group of viruses could have a broader host range than reptiles alone. In addition, there is evidence that this virus, along with other members of this genus – including Charleville virus, which was also isolated in Australia from *Gehyra* (the same host genus as the virus detected here) and sandflies – are arthropod-borne (McAllister et al. 2014; Vasilakis et al. 2019). The lizard *Iridoviridae* found was most closely related to an amphibian and fish virus, with a relatively short genetic distance between them. This is perhaps unsurprising as recent host jumping seems to be commonplace within the *Iridoviridae*; some members of this family can infect fish, reptiles and amphibians (Brenes et al. 2014), and some invertebrate-infecting *Iridoviridae* can also apparently infect vertebrates (Papp and Marschang 2019). In sum, this work supports a model of frequent host-jumping throughout the evolutionary history of most virus families on a backdrop of virus-host co-divergence (Geoghegan, Duchêne and Holmes 2017; Geoghegan and Holmes 2017; Shi et al. 2018).

Our phylogenetic analyses also revealed some geospatial clustering. For virus families with multiple detections, there is little overlap of virus species from different environments. This suggests that lizard viruses are evolving within their host populations in a relatively isolated manner, as might be expected given the generally strong phylogeographic structuring across species ranges (Moritz et al. 2016; Moritz et al. 2018; Potter et al. 2018). It is also of interest that phylogenetic groups that contain viruses sampled from both mesic and arid locations tend to have mesic associated viruses in the basal positions. The hepacivirus phylogeny is particularly enlightening in this respect and it suggests the movement of viruses from one environment to another: both clades of *G. nana* hepaciviruses have viruses collected from mesic environments located in basal phylogenetic positions to those viruses sampled from animals in arid environments. This is consistent with the theory that Australia’s mesic terrestrial biota is mostly ancestral (Byrne et al. 2011). Additionally, more consistent mesic conditions may favor the retention of viral diversity, such that younger taxa dominate in the more climatically variable arid regions of the Kimberley. This could also be associated with the more variable demographic histories of the drier Kimberley compared to the mesic Top End populations (Potter et al. 2018).

The hepacivirus phylogeny also reflects host historical biogeography. As >90% of Australian reptiles are endemic (Chapman 2009), it is expected that viruses infecting Australian reptiles would have minimal opportunities to spread between countries. Indeed, even the currently limited sampling of lizard viruses demonstrates little movement between countries. However, in one clade of hepaciviruses, viruses from Australian *O. marmorata* hosts were separated from viruses from Australian *G. nana* by viruses sampled from China, with the Chinese sampled viruses falling as sister taxa to the Australian *G. nana* viruses. Interestingly, *O. marmorata* are thought to have Gondwanan origins, while the *G. nana* are believed to have immigrated from Asia around the Eocene-Oligocene transition (Oliver and Hugall 2017).

Thus, this clade of hepaciviruses could reflect the historical biogeography of their hosts, with evidence of a lineage of viruses introduced to Australia within immigrant ancestors. As the evolutionary histories of the viruses studied here frequently aligned with the biogeographic and paleoclimatic histories of the hosts, this supports the role of virus evolution in informing animal host ecological histories where sampling is strategic and sufficiently dense (Wilfert and Jiggins 2014). Clearly, however, increased sampling across taxa and biogeographic regions would be useful in confirming this hypothesis.

Although the disease association, if any, of the viruses collected here is unknown, we did identify viruses related to known pathogens, including an *Orthobornavirus*, multiple *Hepaciviruses*, and a member of the *Iridoviridae*. It is noteworthy that seemingly healthy lizards can carry a very high abundance and richness of viruses in the liver, particularly the *Gehyra* species (high viral richness) and *H. planiceps* (high abundance). Viral emergence can have devastating effects on reptile species, such as the Bellinger River snapping turtle that is now critically endangered following mass mortalities due to a novel nidovirus outbreak (Zhang et al. 2018). Although it is unclear where the novel nidovirus (with kidney tropism) originated, the closest known viral relatives were observed in pythons and lizards (usually associated with respiratory disease) (Zhang et al. 2018). Given that lizards make up 59% of non-avian reptiles (Pincheira-Donoso et al. 2013), and one fifth of reptile species are threatened (Cox et al. 2022), this expansion of the lizard virosphere and elucidation of viral genomic sequences may help inform and prepare for potential threats to reptile species.

While this work is the first structured examination of the lizard virome, the sample size is relatively small and there are several caveats. Previous studies have found variation in viromes depending on the age of the individuals sampled (Bergner et al. 2020) and the time of year that sampling occurred (Raghwani et al. 2022). These variables were not controlled in this study. In addition, a larger sample size would enable better comparison between matched species sampled from different environments and a greater sample across the squamate phylogeny would expand these results. The viral families detected here are reflective of the fact that liver samples were sequenced, and as viromes commonly differ by the tissue of sampling, studying a wider range of tissues would be beneficial. Despite these limitations, this study demonstrates that lizards carry a large diversity of viruses, often in high abundance and potentially species-specific. As such they are not only interesting models of vertebrate evolution and ecology, but also serve as good host models by which to study virus ecology and evolution. This work further demonstrates that virus evolution may be a useful tool for understanding or corroborating host biogeographic and paleoclimatic histories.

## Data availability

All raw data (fastq files) generated for this study are available in the NCBI SRA database under BioProject XXXX, BioSample accessions XXXX. Consensus sequences of assembled viral contigs presented in phylogenies are available in GenBank under accession numbers XXXXX-XXXX.

## Acknowledgements

We acknowledge the logistical support provided by Niccy Aitken and Leo Tedeschi at Australian National University while completing the wet-lab component of this study. We acknowledge the Sydney Informatics Hub and the University of Sydney’s high-performance computing cluster Artemis for providing the high-performance computing resources that contributed to the research results reported within this paper.

