## Supplementary Information for "The diverse viromes of Australian lizards are shaped by host taxonomy and habitat"

**Tables S1.** Details of sampled lizard livers in each library (separate excel file).

**Table S2.** Viral data sets used in the phylogenetic analyses.

| *Group-name* | *Figure No.* | *No. taxa* | *Alignment length before trimAl* | *Alignment length used for tree (post trimAl)* |
|---|---|---|---|---|
| *Amnoonviridae* | 4 | 14 | 615 | 512 |
| *Arenaviridae* | 4 | 74 | 1,784 | 1,319 |
| *Astroviridae* | 3 | 91 | 574 | 525 |
| *Bornaviridae* | 4 | 25 | 963 | 947 |
| *Caliciviridae* | 3 | 66 | 598 | 568 |
| *Flaviviridae* | 3 | 231 | 913 | 502 |
| *Hepacivirus only* | 3 | 88 | 608 | 554 |
| *Iridoviridae* | 5 | 27 | 523 | 451 |
| *Picornaviridae* | 3 | 177 | 656 | 479 |
| *Rhabdoviridae* | 4 | 230 | 1,422 | 774 |

**Table S3.** Details of assembled viral nucleotide consensus sequences (likely vertebrate-infecting viruses) (separate excel file).

### Supplementary Figures

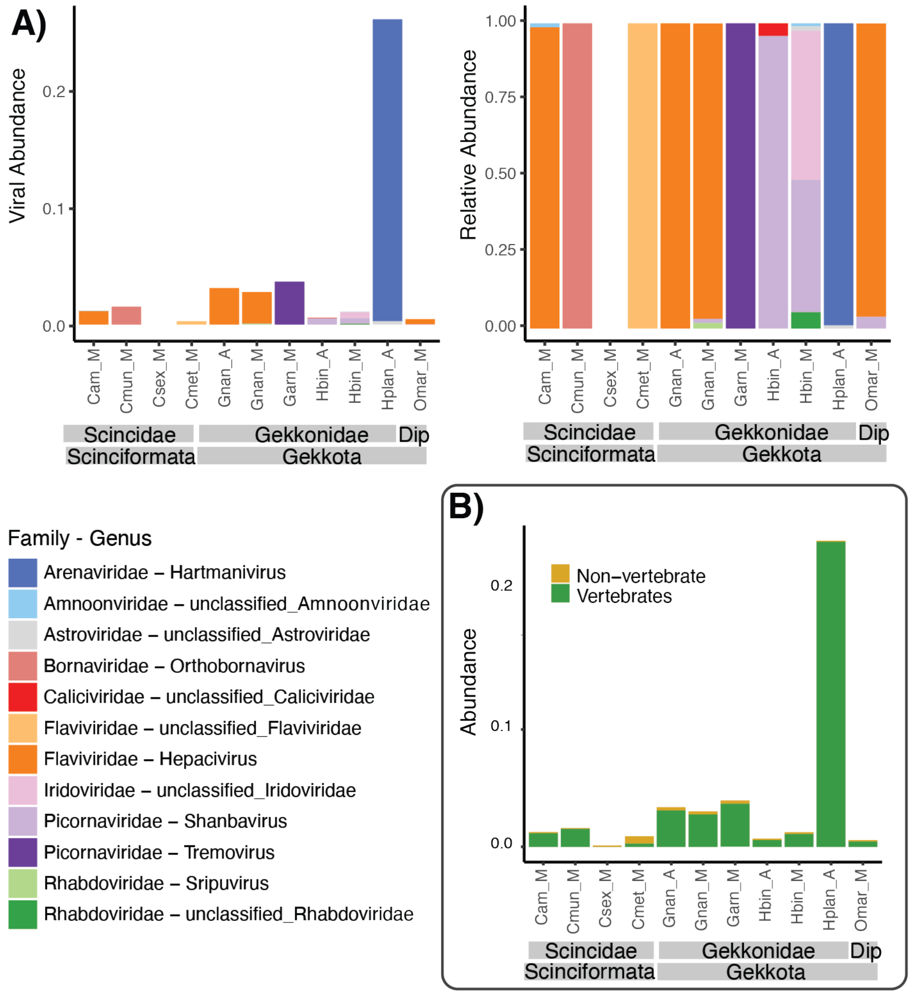


**Figure S1**. **Viral abundance across libraries.** (A) Abundance (left) and relative abundance (right) of likely vertebrate-infecting viruses classified at the virus genus level in each library. Plots classified at the virus family level are presented in Figure 1. (B) Abundance of likely vertebrate-infecting (i.e., biologically relevant) viruses (green) compared to viruses likely infecting the microbiome or potential environmental contaminants (i.e., non-vertebrate viruses; yellow). Library names are as follows: Cam_M, *Carlia amax (collected from a mesic environment)*; Cmun_M, *Carlia munda* (mesic), Csex_M, *Carlia sexdentata* (mesic); Cmet_M, *Cryptoblepharus metallicus* (mesic); Gnan_A, *Gehyra nana* (arid); Gnan_M *Gehyra nana* (mesic); Garn_M, *Gehyra arnhemica* (mesic); Hbin_A, *Heteronotia binoei* (arid); Hbin_M, *Heteronotia binoei* (mesic); Hplan_A, *Heteronotia planiceps* (arid); Omar_M, *Oedura marmorata* (mesic); and are plotted taxonomically, with host order and family names in grey bars (Dip=Diplodactylidae).

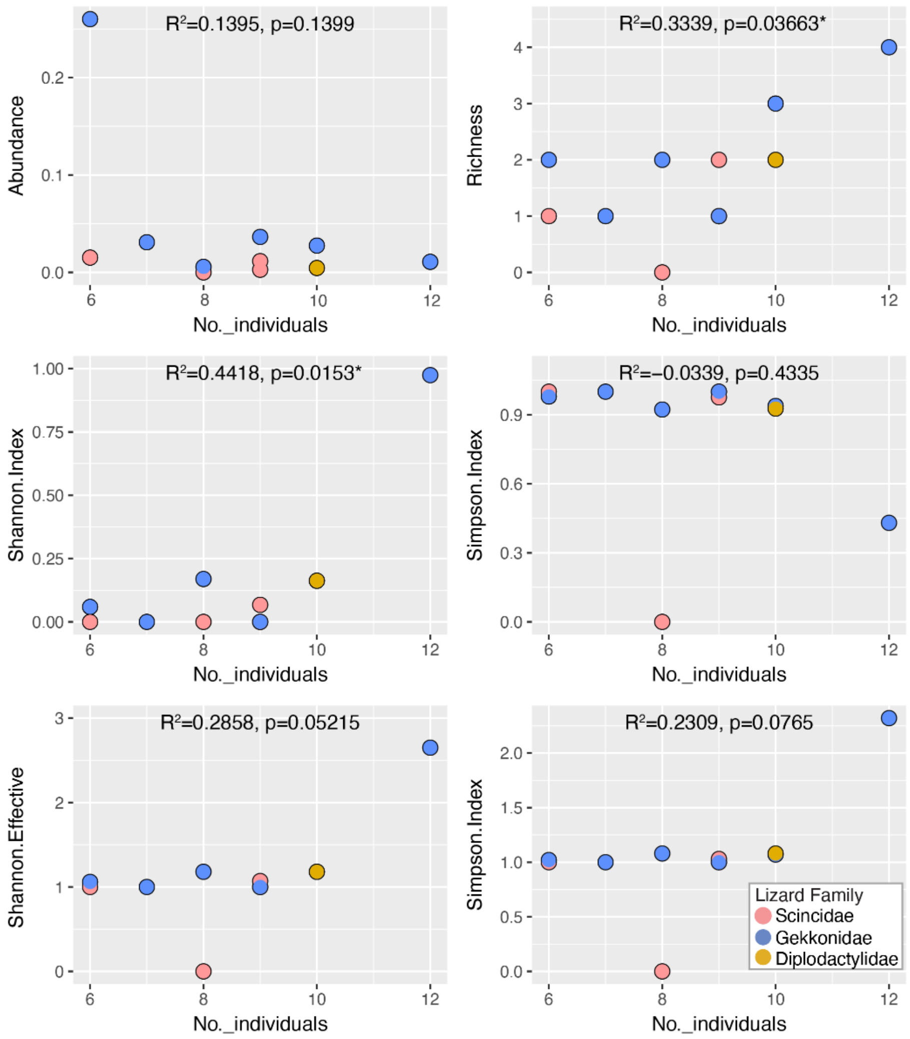


**Figure S2**. **Significant correlation between the number of individuals in each library and viral richness and Shannon Index.** Points are coloured by host family. P-values <0.05 were considered significant and are indicated with an asterisk.

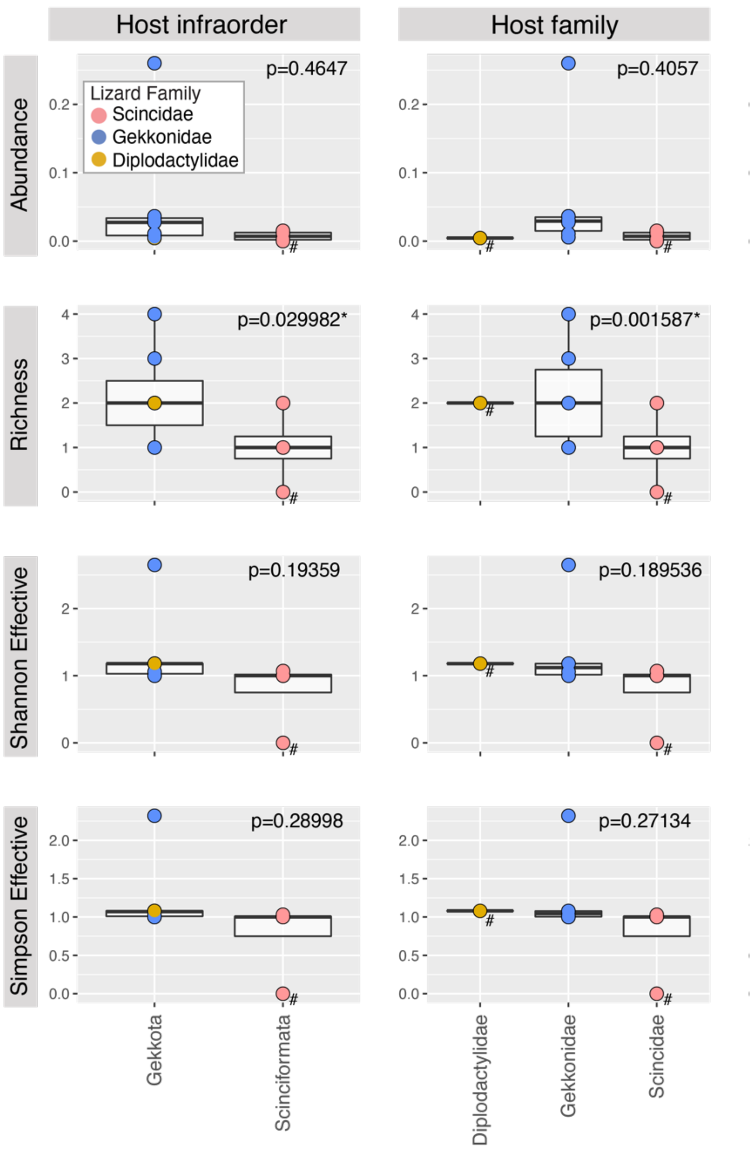


**Figure S3. Viral abundance and alpha diversity according to host taxonomy.** Points are coloured by host family. P-values for host taxonomy in best fit models are specified. P-values <0.05 were considered significant and are indicated with an asterisk. Csex_M and host taxonomic levels comprising only a single library are included in plots for visualisation only and were not included in the final statistical models (denoted by a #).

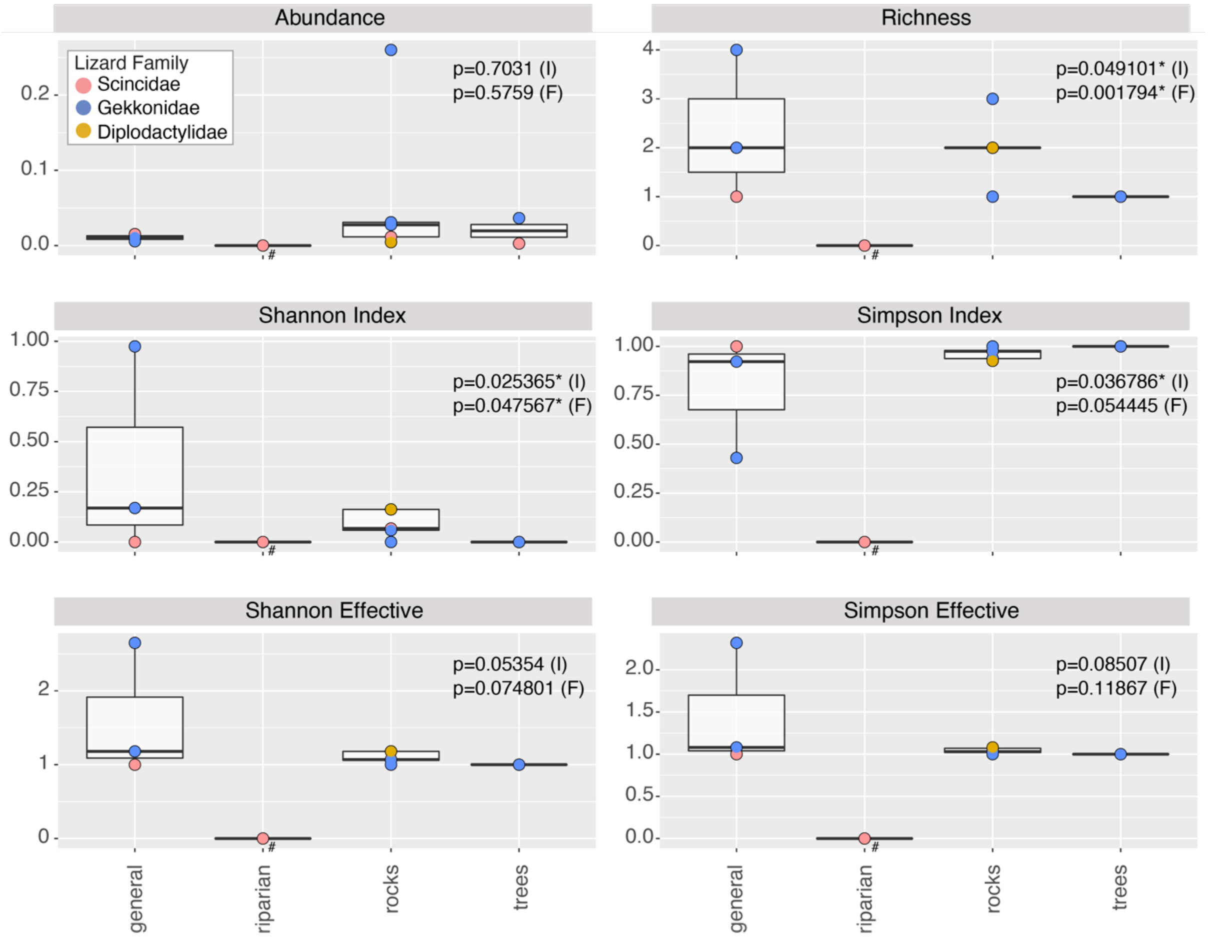


**Figure S4. Viral abundance and alpha diversity by host habitat.** Points are coloured by host family, here as an indicator for host taxonomy (selected for aesthetic reasons only). The Csex_M library (riparian) was included in plots for visualisation only and was not included in the final statistical models (denoted by a #). Additionally, the single Diplodactylidae library, Omar_M (indicated in yellow) was not included in the final statistical models when considering host family taxonomy. P-values for habitat when considered with taxonomy at the level of host infraorder are indicated with an "I", while those considered with taxonomy at the level of host family are indicated with an "F". P-values <0.05 were considered significant and are indicated with an asterisk.

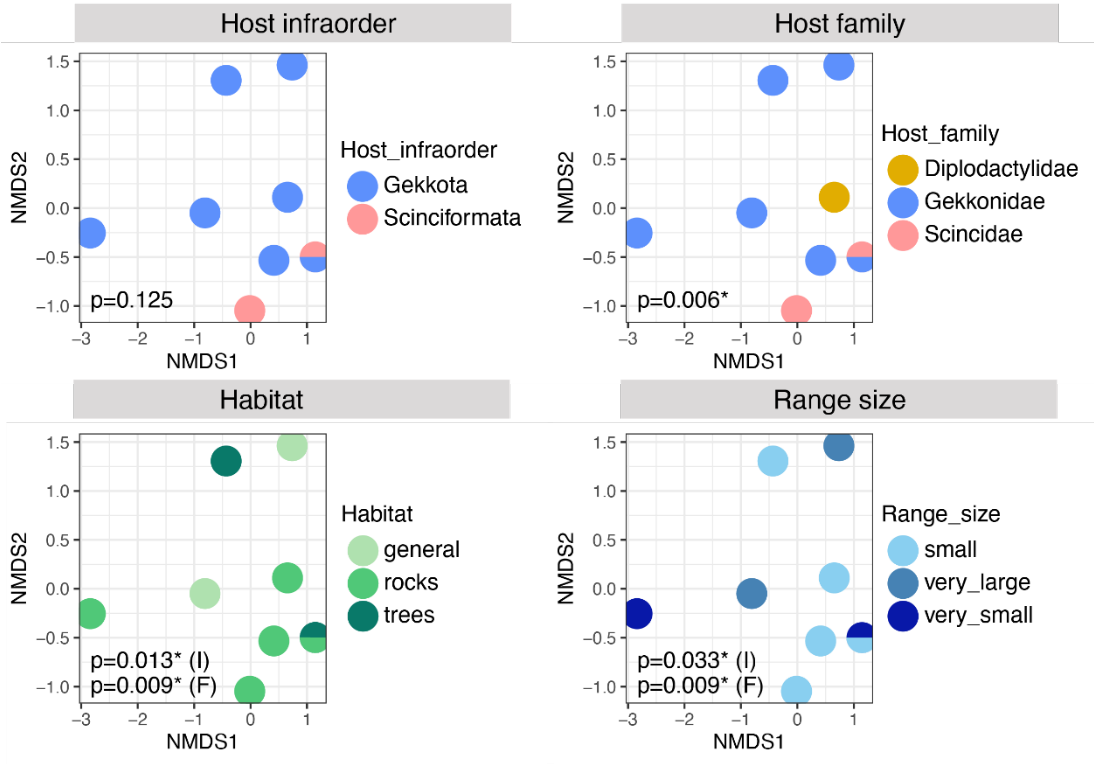


**Figure S5. Beta diversity of viruses in different hosts, habitat and range size.** NMDS plots coloured by considered factors – host infraorder, host family, host habitat and host range size. Overlapping points are depicted by a half circle. P-values indicated on habitat and range size plots are both those calculated in models considering the host family level ("F") as well as those calculated considering the infraorder level ("I"). P-values <0.05 were considered significant and are indicated with an asterisk. Csex_M was not included as no biologically relevant viruses (i.e., vertebrate-infecting viruses) were found in this library.
